# Adolescent substance use and functional connectivity between the ventral striatum and hippocampus

**DOI:** 10.1101/847749

**Authors:** Edward D. Huntley, Hilary A. Marusak, Sarah E. Berman, Clara G. Zundel, Joshua R.B. Hatfield, Daniel P. Keating, Christine A. Rabinak

**Author notes:** Corresponding Author: Hilary Marusak, PhD, Department of Psychiatry & Behavioral, Neurosciences, School of Medicine, Wayne, State University, Address: 3901 Chrysler Service Dr., Suite, 2B, Detroit, MI, 49201. Authors contributed equally to the work (Edward D. Huntley, Hilary A. Marusak).

## Abstract

Neurodevelopmental explanations for adolescent substance use have focused on heightened sensitivity of the brain’s reward system, centered around the ventral striatum (VS). Recent evidence demonstrates increased functional connectivity between the VS and hippocampus in adolescents relative to adults, suggesting that the adolescent brain may learn from subsequent exposure to risks/rewards. However, a link between VS-hippocampal circuitry and adolescent substance use has yet to be established. Two separate longitudinal studies were conducted to evaluate whether variation in VS-hippocampal resting-state functional connectivity (rs-FC) predicts subsequent adolescent substance use. Study 1 consisted of 19 youth recruited from a high sociodemographic risk population (N = 19; 14 female; 47% Black Non-Hispanic, 32% White Non-Hispanic). To replicate results of Study 1, Study 2 utilized data from the National Consortium on Adolescent Neurodevelopment and Alcohol, an ongoing multi-site imaging study (N= 644; 339 female; 11% Black Non-Hispanic, 11% Hispanic/Latino, 66% White Non-Hispanic). Resting-state fMRI data were collected at a baseline time point and lifetime and past year self-reported substance use was collected at a follow up visit. Regression models tested whether baseline VS-hippocampal rs-FC predicted substance use at follow up. Across both studies, higher VS-hippocampal rs-FC at baseline predicted greater substance use at follow up. These data provide the first evidence linking increased VS-hippocampal connectivity with greater adolescent substance use. Results fit with the emerging idea that adolescent substance use is driven by not only a heightened sensitivity to reward, but also a stronger link between reinforcement learning and episodic memory for rewarding outcomes.

## Introduction

Experimentation with alcohol, marijuana, and other substances is common among adolescents (1, 2). However, adolescent substance use is associated with increased risk of substance use disorders in adulthood, which have a lifetime prevalence of approximately 10% in the US (3). Adolescent substance use has also been linked to later cognitive problems (e.g., learning, memory, attention), which may relate to changes in adolescent brain structure and function (4, 5). Adolescent substance use tends to co-occur with engagement in other health risk behaviors, including sexual activity, violent behavior, and reckless driving (6). Together, the consequences of substance use for a substantial number of adolescents are severe, including lifelong health problems and accidents and unintentional injuries, which represent the leading cause of death among young people ages 10-24 in the US (7). The enormous public health costs of adolescent substance use (8, 9) have motivated research into the underlying neurobiological mechanisms, and to identifying early markers that predict subsequent engagement.

A propensity for substance use and other risk-taking behavior during adolescence is thought to arise from heightened sensitivity of the brain’s reward circuitry (10, 11). Prior neuroimaging studies indicate that key areas of reward circuitry, particularly the ventral striatum (VS), are more responsive to incentive-based stimuli (e.g., peer approval) in adolescents relative to both adults and children (12, 13). The VS plays a central role in reward-seeking behavior and learning, and is a key target of dopaminergic projections from midbrain regions (14). Given that midbrain dopaminergic systems are involved in reward-seeking and incentive motivation (15, 16), heightened VS reactivity during adolescence is thought to convey an increased sensitivity to positive outcomes, even if the behavior is risky (17). In line with this notion, higher VS reactivity has been shown to correspond with more risky decisions during risk taking paradigms (18, 19), and with adolescent substance use (20).

Recent evidence suggests that it is not only a heightened neural sensitivity to rewarding outcomes that drives risk-taking behavior during adolescence, but also an increased ability to *learn* from positive reinforcement (21). Recent work by Davidow and colleagues (22) demonstrated that, relative to adults, adolescents show better reinforcement learning and a stronger link between reinforcement learning and episodic memory for rewarding outcomes. This behavioral effect was associated with stronger functional connectivity (FC) between the striatum and the hippocampus during reward learning, which may reflect greater integration of reward-related information into long-term memory (22). Together, the VS, hippocampus, and midbrain form a critical reward-memory loop that signals the motivational significance of events and modulates episodic memory formation in the hippocampus (23, 24). Connections within VS-hippocampal circuitry are therefore critically involved in the strengthening of reward-guided habits, actions, and episodic memory for motivational events, thereby allowing past experiences to adaptively guide behavior (23, 24). In light of the recent findings of Davidow et al. (22), VS-hippocampal circuitry may be involved in the propensity for substance use during adolescence. However, a direct link between VS-hippocampal circuitry and adolescent substance use has yet to be established.

The present study was designed to test the hypothesis that increased FC between the VS and hippocampus predicts greater subsequent substance use in adolescence. This hypothesis is based on the notion that youth who show greater VS-hippocampal FC may be sensitized to better learn from rewarding experiences and may therefore be more likely to subsequently apply this learning and engage in substance use (21, 22). Thus, here we examine the brain as a potential predictor of future behavior. We test this hypothesis first with a longitudinal imaging study (Study 1) performed in a sample of youth at high sociodemographic risk for substance use and associated negative outcomes (25) (Detroit, Michigan, USA). We then performed an independent replication study in a large, publicly available dataset (National Consortium on Adolescent Neurodevelopment and Alcohol, NCANDA), using longitudinal data collected in n = 644 US adolescents and young adults (Study 2).

## Materials and Methods

Two separate longitudinal studies were conducted (Studies 1 and 2) to examine brain patterns predicting subsequent substance use during the high-risk period of adolescence. All included participants had resting-state functional magnetic resonance imaging (fMRI) scans at time point 1 (T1) and returned for a subsequent substance use assessment during adolescence (time point 2, T2).

### Participants

#### Study 1

Nineteen youth, predominantly female (*n =* 14), participated in this study at Wayne State University (WSU; see Table-I and Figure-1). The age range for T2 (15-17 years) was selected to roughly coincide with a large-scale international study (>5,000 individuals) showing that sensation-seeking increases during adolescence, peaks around age 19, and subsequently declines into adulthood (26). Self-regulation also increases during adolescence, and reaches a plateau between the ages of 23 and 26 (26). These observations are congruent with other age gradients in risk-taking behavior, such as arrests for criminal behavior and substance use (1). Participants were recruited through advertisements posted on the WSU website, Craigslist (Detroit, Michigan), printed flyers, or through area behavioral medicine clinics. Exclusionary criteria included lack of parental involvement, lack of English proficiency, history of brain injury, neurological or movement disorders, or presence of MRI contraindication.

**Figure 1.**
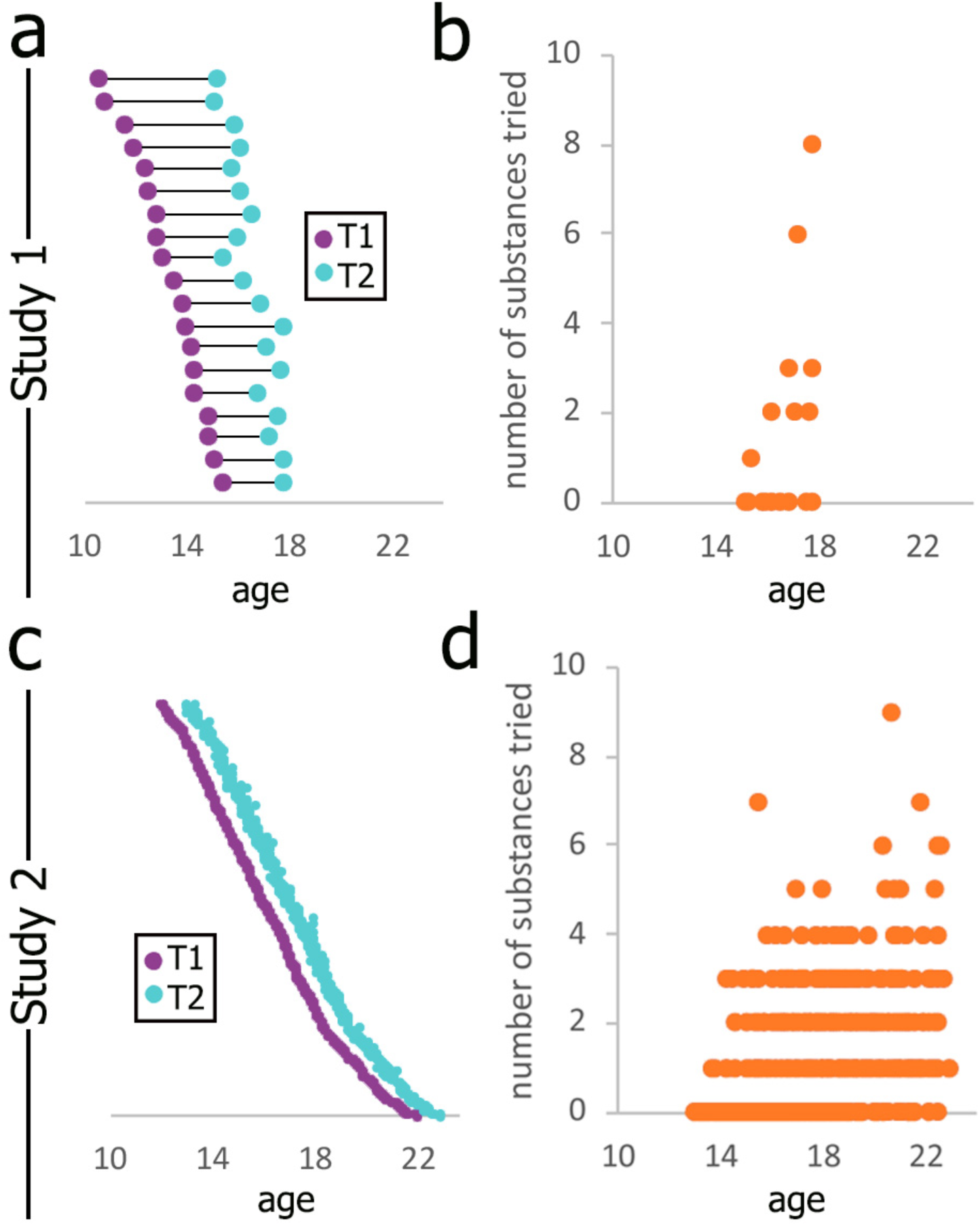
Age at baseline (T1) and follow up (T2) visits, and substance use endorsed at T2, by age, for Study 1 (top) and Study 2 (bottom). MRI scanning occurred at T1. The substance use survey occurred at T2 (and also T1 for Study 2). Abbreviations: T1, Time 1; T2, Time 2; MRI, magnetic resonance imaging. See also Tables 1 and 2.

Consistent with the study location (Detroit, Michigan), the sample was racially and economically diverse (Table1). Research shows that minority, lower-income populations are at increased risk for engaging in health risk behaviors during adolescence, and are more likely to suffer from associated negative consequences (e.g., substance use disorders (27-30)). The follow up visit (T2) occurred 2-4 years after time point 1 (*M* = 3.26, *SD* = 0.75).

#### Study 2

Study 2 reports on data collected from 644 adolescents and young adults from NCANDA, a large multi-site study that aims to examine the neurocognitive correlates of alcohol use in a nationally representative (sex and racial/ethnic groups) sample of US adolescents (31). To replicate results of Study 1, we included participants with resting-state fMRI data collected at T1 and who reported on substance use at the 1-year follow up visit (T2; see Figure-1C; 1 ± .12 years between visits). To identify predictors of future substance use risk, youth were recruited for the NCANDA study for limited substance use at T1 but oversampled youth at high risk (e.g., family history of substance use disorder). NCANDA participants were recruited across five sites: Duke University, University of Pittsburgh Medical Center, Oregon Health & Science University, University of California, San Diego, and SRI International. Exclusionary criteria included lack of parental involvement, lack of English proficiency, psychotropic medication use, early developmental problems, psychiatric disorders, pervasive developmental disorder, or MRI contraindications (see (31) for full list). Similar to Study 1, the Study 2 sample was racially and economically diverse (Table 1).

**Table 1.**
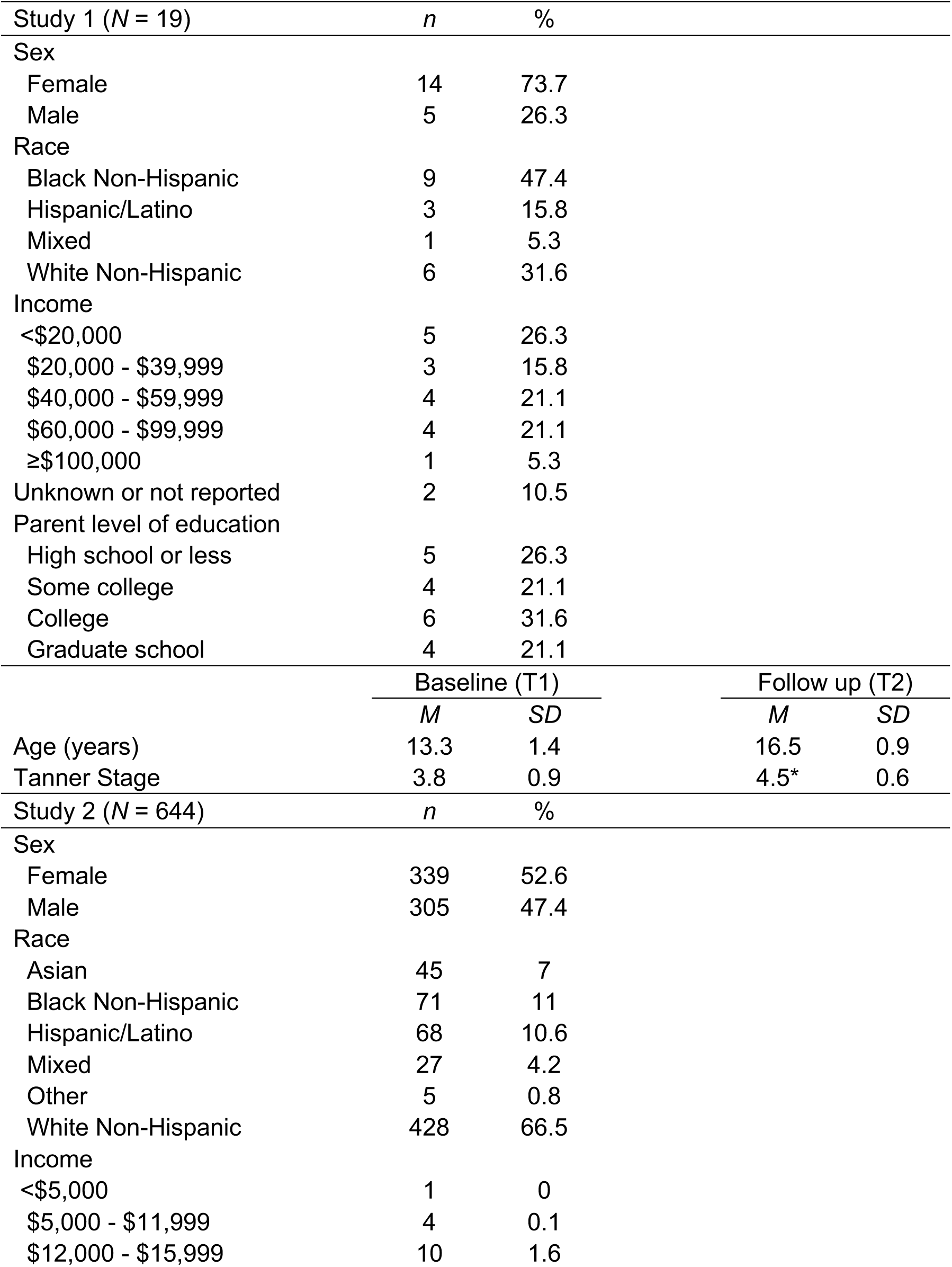
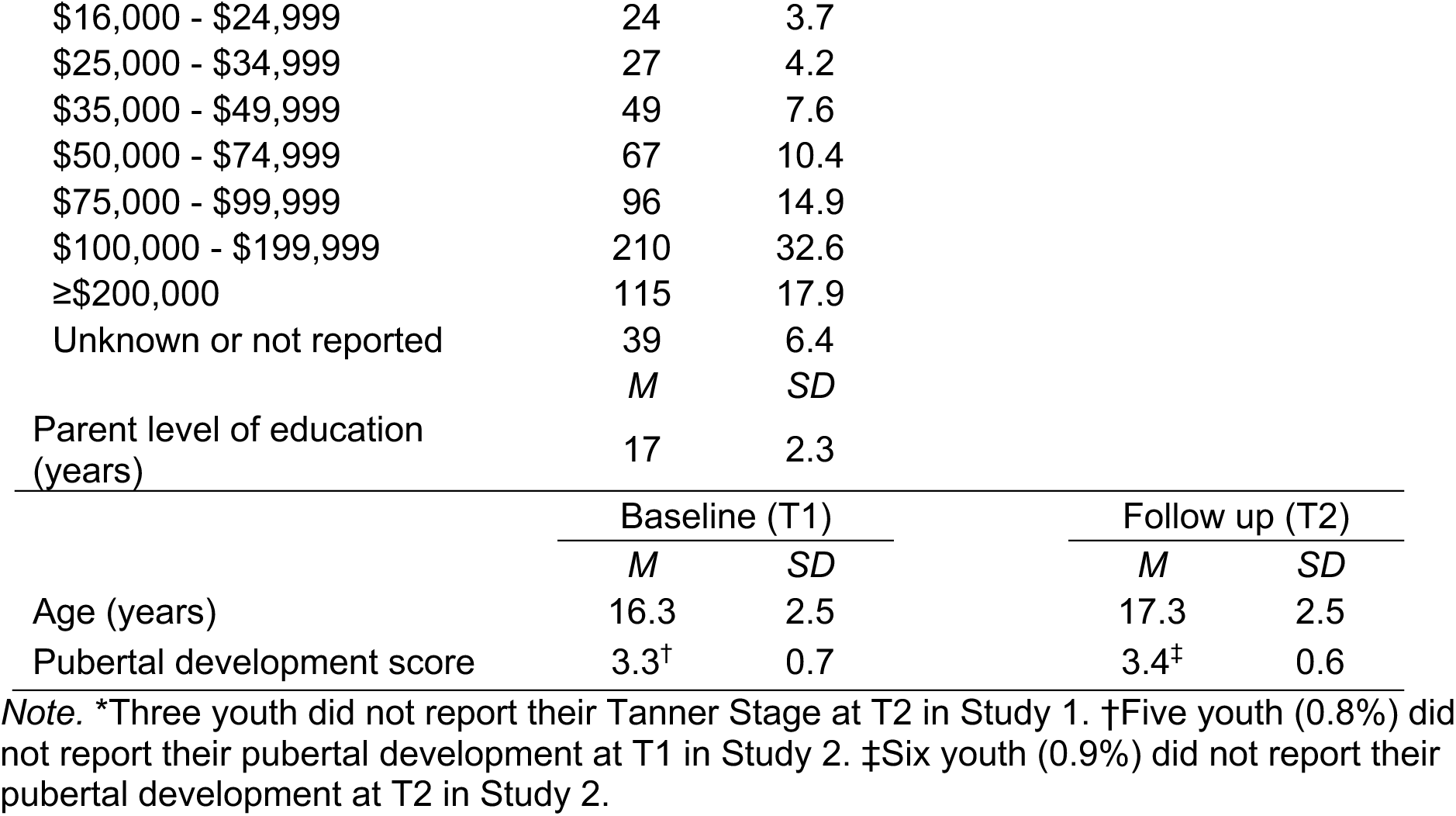
Demographic data for each study

#### Both studies

All participants provided written informed/parental informed consent and child or adolescent assent prior to participation. Study procedures were approved by the local IRB and have been performed in accordance with the ethical standards laid down in the 1964 Declaration of Helsinki and its later amendments. For both studies, a Certificate of Confidentiality issued by the US Department of Health and Human Services was obtained to help further protect the privacy of participants in disclosing substance use.

### Substance use

Adolescent self-report of lifetime and past year substance use was evaluated at T2 using a summary score, by dichotomizing responses for engagement in various substance use categories (alcohol, cigarettes, marijuana, amphetamines, narcotics, sedatives, hallucinogens, cocaine, inhalants, opiates, illicit use of prescription drugs). In particular, those who indicated “0 occasions” were coded as not using this substance and those who reported any occasions were coded as using this substance. Of note, the number of substance use categories tried was positively correlated with the number of episodes of alcohol or drug intoxication (*r* = .48, *p* < .001). Neuroimaging results for number of episodes were consistent with reported results. See Supplemental Material for further information.

### Imaging acquisition and preprocessing

See Supplemental Material for details on collection of resting-state fMRI data and approaches to reduce the potential impact of head motion on results. FC pre-processing and group-level analyses were conducted using identical procedures across both studies. In brief, fMRI preprocessing was conducted using SPM8 software (Statistical Parametric Mapping; http://www.fil.ion.ucl.ac.uk/spm/) and included slice-timing correction, realignment, spatial normalization to the Montreal Neurological Institute (MNI) template, and smoothing using a 6-mm Gaussian kernel.

### VS seed-based resting-state functional connectivity analysis

Resting-state functional connectivity (rs-FC) was measured using a seed-based approach. Left and right VS seeds were defined as spherical regions of interest (ROIs; 4-mm radii; 33 voxels) centered at x = ±10, y = 14, z = 0, following a previous rs-FC study in youth (32). Rs-FC of left and right VS seeds with the rest of the brain was determined using Pearson bivariate correlation with the CONN Toolbox (ver.12.p; www.nitrc.org/projects/conn). Resulting correlation values were normalized using Fisher z-transformation.

### Group-level VS rs-FC analysis

Study 1 and Study 2 were analyzed separately. First, to establish the basic network topology and compare this to previous research, we evaluated rs-FC of left and right VS across each sample, using one-sample *t*-tests in SPM8. A liberal threshold of *p* < .005, 10-voxel minimum was used to visualize network topology, and areas surviving whole brain cluster-level correction (Study 1: *p* < .001, 118-voxel minimum; Study 2: *p* < .001, 550-voxel minimum) are indicated in Tables-S2 and S3. The whole brain threshold was determined by computing the spatial autocorrelation of the data via AFNI’s 3dFWHMx and subsequently performing Monte Carlo simulations (10,000 iterations) using 3dClustSim (compile date July 22, 2016; National Institute of Mental Health, Bethesda, MD; https://afni.nimh.nih.gov). Our main analysis tested whether VS-hippocampus connectivity at T1 predicted substance use reported at T2, using a regression analysis in SPM8, with lifetime number of substances tried as the variable of interest and age at T1 as a variable of no interest. Additionally, given the variability in length of time between visits in Study 1, months between T1 and T2 were included as an additional variable of no interest in Study 1 analyses. We performed follow-up analyses to test whether lifetime number of substances tried at T1 also predicted VS-hippocampal connectivity at T1 (Study 2 only; substance use data unavailable for T1 in Study 1). Results were considered significant within an *a priori* hippocampus ROI, based on Davidow et al.(22) (*x* = −16, *y =* −8, *z =* −20, 12-mm radius sphere), using small-volume familywise error correction, *pFWE* ≤ .05. A functional ROI was created around peak effects from Study 1 for use in Study 2 (6-mm radius sphere). A complementary whole brain analysis was performed, and effects were considered significant at the whole brain corrected threshold (see above). All coordinates provided in this report are given in MNI convention. Given evidence of positive skew in substance use scores, we confirmed that results reported did not change when applying a square root transformation, which resulted in a more normal distribution. Observed results remained significant when using the transformed variable.

## Results

### Substance use across Study 1 and 2 samples

42% and 48% of participants in Study 1 and Study 2, respectively endorsed lifetime use of at least one substance at T2, with alcohol, marijuana, and cigarettes being the most common (see Table-II for summary of lifetime and past year use). Lifetime and past year substance use was highly interrelated across both studies (*r*s >.9, *p*s < .001). Across both studies, older youth reported more substance use than younger youth (Study 1: *r* = .5, *p* = .03; Study 2: *r* = .45, *p* < .001; see Figure-1b and 1d, respectively). Overall, these patterns are consistent with previous reports (1, 2)

**Table 2.** Summary of lifetime adolescent substance use for each study

| Study 1 (N = 19) | Lifetime |  | Past 12 mo. |  |  |  |
| --- | --- | --- | --- | --- | --- | --- |
|  | *n* | % |  | *n* | % |  |
| Alcohol | 7 | 36.8 |  | 7 | 36.8 |  |
| Cigarettes | 4 | 21.1 |  | 4 | 21.1 |  |
| Marijuana | 6 | 31.6 |  | 5 | 26.3 |  |
| Amphetamine | 1 | 5.3 |  | 1 | 5.3 |  |
| Narcotic | 1 | 5.3 |  | 1 | 5.3 |  |
| Sedatives | 2 | 10.5 |  | 2 | 10.5 |  |
| Other illicit drugs | 1 | 5.3 |  | 1 | 5.3 |  |
|  | *M* | *SD* | Min -<br>Max | *M* | *SD* | Min -<br>Max |
| Substance exposure | 1.3 | 2.2 | 0 - 8 | 1.2 | 2.2 | 0 - 8 |
| Study 2 (N = 644) | Lifetime |  | Past 12 mo. |  |  |  |
|  | *n* | % |  | *n* | % |  |
| Alcohol | 281 | 43.6 |  | 260 | 40.4 |  |
| Cigarettes | 107 | 16.6 |  | 74 | 11.5 |  |
| Marijuana | 188 | 29.5 |  | 159 | 24.7 |  |
| Amphetamine | 12 | 1.9 |  | 8 | 1.2 |  |
| Narcotic | 6 | 0.9 |  | 3 | 0.5 |  |
| Sedatives | 0 | 0 |  | 0 | 0 |  |
| Other illicit drugs | 33 | 5.1 |  | 24 | 3.7 |  |
|  | *M* | *SD* | Min -<br>Max | *M* | *SD* | Min -<br>Max |
| Substance exposure | 1 | 1.3 | 0 - 9 | 0.8 | 1.2 | 0 - 6 |

### VS rs-FC across Study 1 and 2 samples

Similar patterns of left and right VS rs-FC were observed across both samples (Figure 2 and Tables S2-S3), with positive connectivity observed across regions of the orbitofrontal cortex, medial temporal lobe (including hippocampus), medial frontal regions, insula, thalamus, and basal ganglia. Overall, these patterns are similar to those observed in a previous study of VS rs-FC in youth (32).

**Figure 2.**
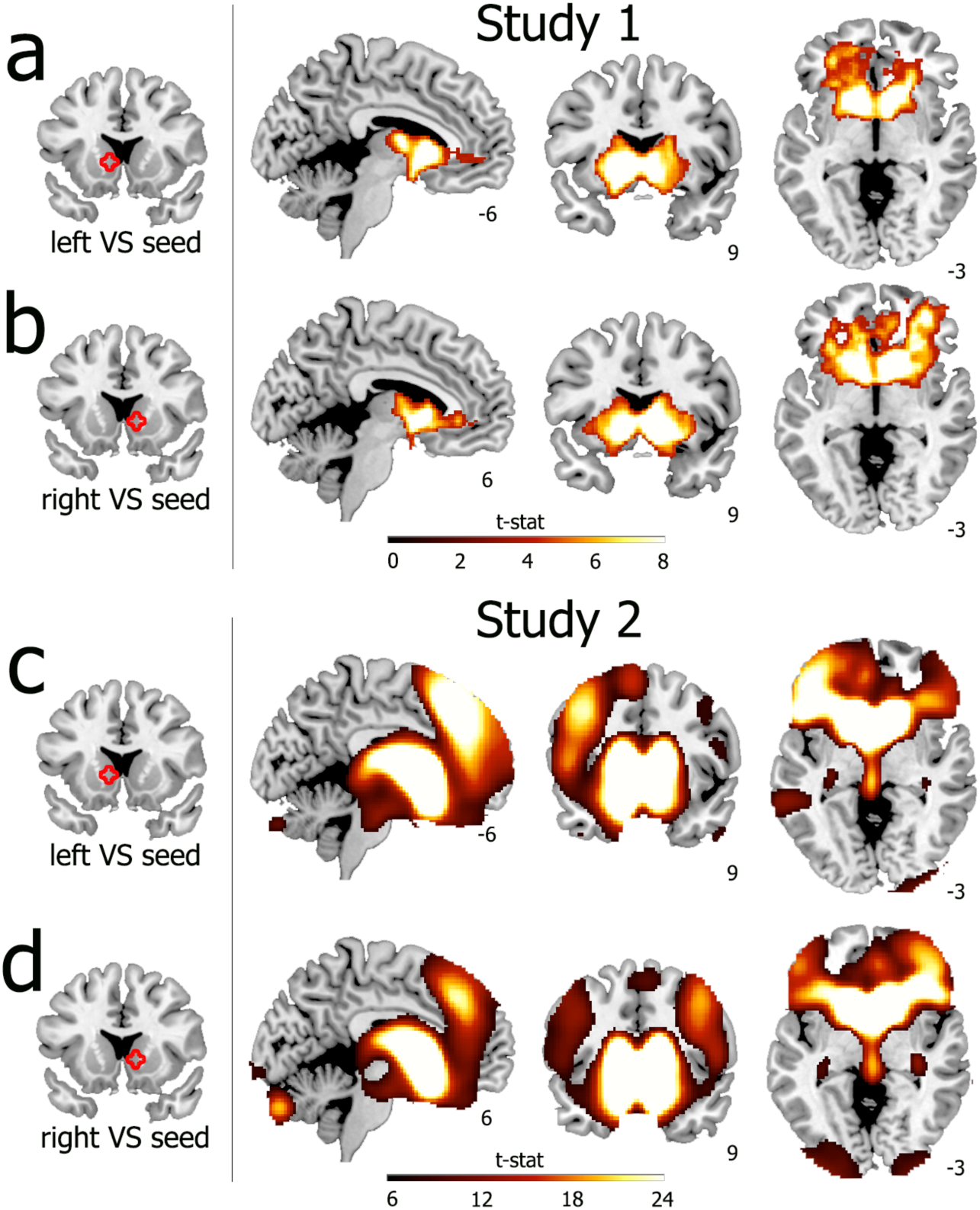
Overall resting-state functional connectivity of left and right VS for Studies 1 (a,b) and 2 (c,d). Left and right VS seed regions of interest are defined based on Porter et al. (32) Connectivity maps shown at the corrected whole-brain threshold (Study 1: *p* < .001, 118-voxel minimum; Study 2: *p* < .001, 550-voxel minimum). Abbreviations: VS, ventral striatum. Connectivity values are given as Fisher’s z-transformed Pearson’s correlations.

### VS-hippocampal connectivity is associated with later substance use

#### Study 1

*A priori* ROI analyses showed that higher right VS rs-FC with the left hippocampus (*pFWE* = .041, *Z* = 3.59, *x* = −20, *y =* −18, *z =* −24, 28 voxels) at T1 was associated with more substance use at T2 (Figure 3). There were no significant effects of substance use with left VS or right hippocampus. Analyses repeated with the addition of motion despiking were consistent with these results (*pFWE* = .023, *Z =* 3.91, x = −20, *y =* −18, *z* = −24, 32 voxels).

**Figure 3.**
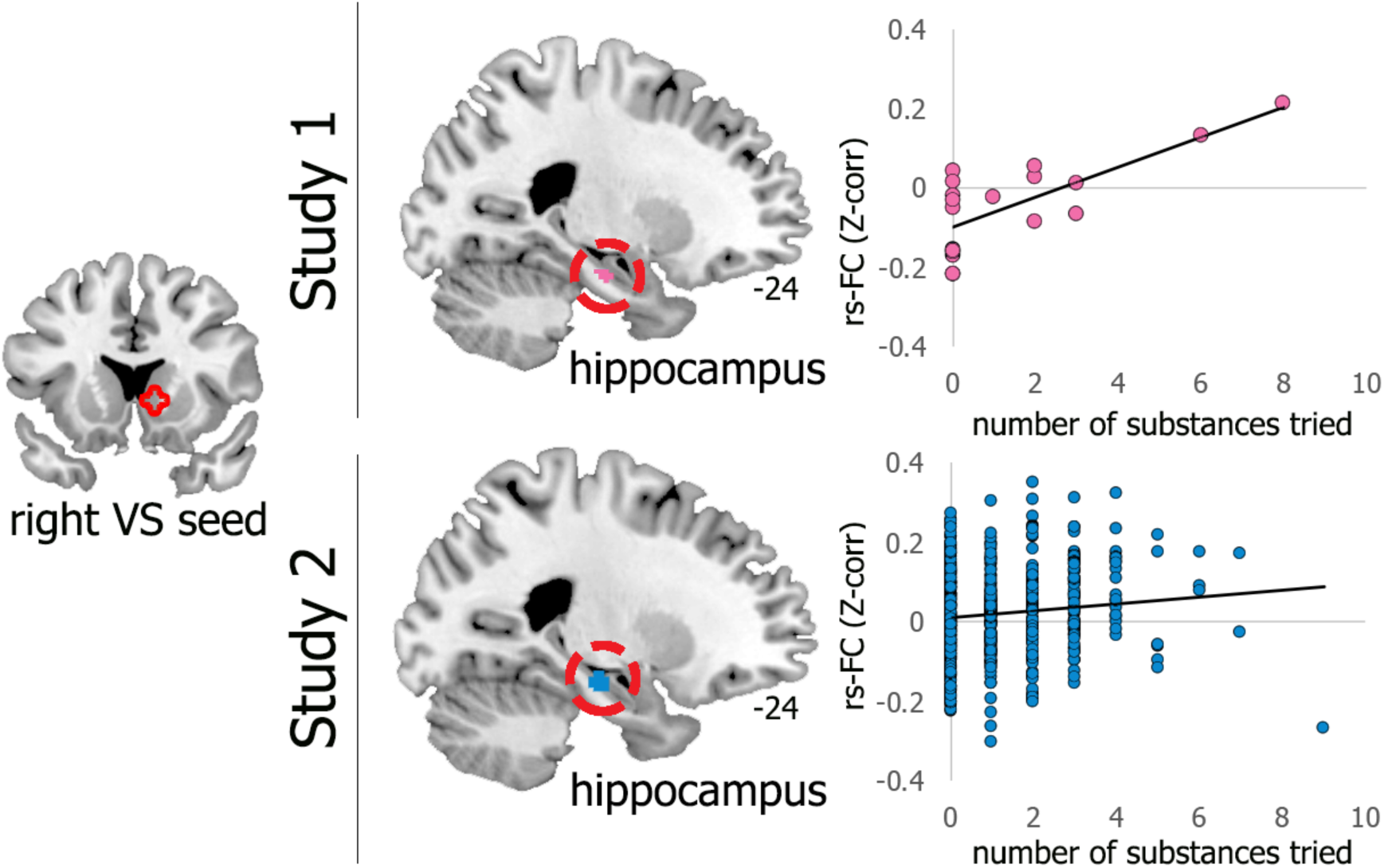
Higher VS functional connectivity with the hippocampus is subsequently associated with a greater number of substances tried during adolescence. Results significant at a small-volume familywise error-corrected threshold of *pFWE* < .05, displayed at *p <* .005 masked in anatomical hippocampus (AAL atlas). Right: Rs-FC values (Fisher’s Z-transformed Pearson’s correlations) were extracted for each participant from group peaks in hippocampus and plotted by substance use for display purposes only. Abbreviations: VS, ventral striatum; rs-FC, resting-state functional connectivity.

#### Study 2

Consistent with results of Study 1, higher right VS rs-FC with the left hippocampus (*pFWE* = .048, *Z* = 2.63, *x* = −22, *y =* −16, *z =* −20, 7 voxels) at T1 was associated with more substance use at T2 (Figure 3). There were no significant effects of substance use with left VS or right hippocampus. Analyses repeated with the addition of motion despiking were consistent with these results (*pFWE* = .05, *Z =* 2.61, x = −22, *y =* −16, *z* = −1, 6 voxels). Importantly, there was no association between lifetime substance use reported at T1 and VS-hippocampal rs-FC at T1.

### Whole brain results

#### Study 1

Whole brain analyses indicated one region spanning fusiform gyrus, cerebellum, and inferior occipital lobe (*x =* 24, *y =* −66, *z =* −20, *Z =* 3.89, 135 voxels; Figure S1) showing lower connectivity with left VS in youth who reported higher substance use. No other areas were significant at the corrected whole brain threshold.

#### Study 2

No areas reached significance at the corrected whole brain threshold.

## Discussion

Important recent work by Davidow and colleagues (22) demonstrated increased VS-hippocampal connectivity in adolescents relative to adults, which corresponded with stronger integration of reward-related reinforcement learning into long-term memory. These results provide new evidence that the increased propensity towards substance use in adolescence may be driven not only by a heightened neural sensitivity to reward– the focus of much contemporary research– but also an increased sensitivity to encode reward-based behavior or cues into long-term memory. However, a direct link between VS-hippocampal circuitry and engagement in substance use was not tested in that study. The present study used two independent samples to identify associations between VS-hippocampal connectivity and substance use during adolescence. Across both studies, we found that increased rs-FC within VS-hippocampal circuitry longitudinally predicted a greater number of substances that adolescents reported using.

The VS, hippocampus, and also midbrain regions are key components of a dopaminergic circuitry that signals motivationally important events and modulates the formation of long-term memory for those events (33). These regions form a critical functional loop that provides top-down signals of novelty (arising from the hippocampus) and contributes to the bottom-up formation of long-term memory in the hippocampus. In particular, detection of novel or motivationally salient stimuli (via hippocampus) leads to increased activation of the VS and midbrain regions. Activation of these regions, in turn, causes a release of dopamine and serves to enhance long-term potentiation in the hippocampus (24). Results of Studies 1 and 2 indicate increased integration within VS-hippocampal circuitry in youth subsequently reporting more substance use. These findings provide support for the notion that substance use during adolescence may be driven by dopaminergic sensitivity to motivationally-relevant cues, and/or how strongly these cues imprint into memory. Although this hypothesis should be tested in future research, increased neural connectivity within this core dopaminergic circuitry may reflect a sensitivity such that reward-based behavior or cues are more strongly encoded, and thus, more likely to guide later choices. Higher VS-hippocampal rs-FC during the transition from childhood and adolescence may serve as a marker for increased health risk behavior during adolescence. Together with other critical risk factors (e.g., peers, family factors, genetic background), a neural marker could be of great service for designing neurodevelopmentally-informed early interventions.

Interestingly, VS-hippocampal rs-FC at T1 was associated with substance use reported at T2 (but not T1). This suggests that strength of integration within VS-hippocampal circuitry predicts a tendency to engage in future (but not current) substance use, or the association between substance use and VS-hippocampal circuitry strengthens over time. Indeed, substance experimentation tends to follow a developmental trajectory such that engagement in substance use and other risk behaviors increases rapidly from early-to-mid adolescence, and peaks during mid-to-late adolescence (1, 2, 34).

Across both studies, the included samples were both racially and economically diverse and included a large number of minority adolescents residing in lower income households (Table 1). Research shows urban, minority, and lower-socioeconomic populations are at heightened risk for engaging in health risk behaviors during adolescence, and are more likely to suffer from associated negative consequences (27-30). Understanding risk pathways is thus especially important in high-risk populations, and requires inclusion of more understudied populations in medical (35) and neuroscientific research (36).

### Limitations

Based on prior work by Davidow et al. (22), we took a hypothesis-driven approach and focused on VS-hippocampal circuitry. This approach reduces the potential for spurious findings and limits our focus on dopaminergic circuitry that is known to be critical for reward-driven behavior and memory integration. However, it does not preclude the potential role of other brain regions and connections that may predict later substance use. While our complementary whole brain analyses in Study 1 linked substance use to altered VS connectivity with a region spanning the fusiform gyrus, cerebellum, and inferior occipital lobe, this result did not replicate in Study 2. Future studies should evaluate other brain regions and larger-scale networks (e.g., salience network). Next, substance use was assessed using adolescent self-report. Although the measure shows good reliability and construct validity (see *Methods*), future studies should consider the additional evaluation of an objective measure of adolescent substance use (e.g., hair samples) or related health risk behaviors (e.g., risky driving fMRI task (37)). Furthermore, we measured VS connectivity during resting-state, and therefore patterns observed here may not be consistent with those observed during reward processing or a risk-taking task. However, recent research suggests that patterns observed in the resting-state predict individual differences during behavioral tasks (38). Nevertheless, it will be important to test whether VS-hippocampal connectivity is similarly increased during a risk-taking task as an additional test of the hypothesis. Similarly, our interpretation of these rs-FC data assume a functional process that is previously described in the extant literature (e.g., 24), and is a limitation of the current work. We cannot confirm that the FC observed in these studies is directly related to risk/reward behavior, and future work should incorporate cognitive tasks that engage risk/reward processes to further evaluate our interpretations of VS-hippocampal rs-FC.

### Conclusion

Many health risk behaviors are established during adolescence and frequently maintained into adulthood, influencing lifelong health and wellbeing. Research into neurobiological drivers or predictors of future engagement in substance use and other risk behaviors during adolescence may help to reduce this health burden by informing intervention strategies and governmental policies. Recent research into the neurobiological origins of substance use during adolescence suggests that the adolescent brain is not only more sensitive to reward, but also more sensitive to reinforcement than the adult brain, which may be reflected in the observed increased connectivity between the VS and hippocampus (22). The present study provides the first evidence linking increased VS-hippocampal connectivity with subsequently greater substance use during adolescence. These findings prompt further investigation into role of dopamine-rich VS-hippocampal circuitry in susceptibility to adolescent substance use, and as potential markers for identifying at-risk youth.

## Acknowledgements

The authors would like to thank Shelley Paulisin, Sajah Fakhoury, Allesandra Iadipaolo, Limi Sharif, Xhenis Brahimi, Farah Sheikh, Suzanne Brown, Laura M. Crespo, Kelsey Sala-Hamrick, Klaramari Gellci, Maria Tocco, Andrea Bedway, Angela Vila, Yashwanth Katkuri, Richard Genik, and Pavan Jella of Wayne State University (WSU) for assistance in participant recruitment and data collection. Thank you also to the adolescents and families who generously shared their time to participate in this study.

## Funding

Research reported in this publication was supported, in part, by the Departments of Pharmacy Practice and Pediatrics, and the Merrill Palmer Skillman Institute of WSU. Dr. Rabinak is supported by NIH National Institute of Mental Health grant R61 MH111935. Dr. Marusak is supported by NIH National Institute of Mental Health grant K01 MH119241. Drs. Huntley and Keating are supported by the NIH *Eunice Kennedy Shriver* National Institute of Child and Human Development (R01 HD075806).

## Conflict of interest

The authors declare that they have no conflict of interest.

